# Inhibitory budget matching constrains homeostatic plasticity in competitive spiking networks

**DOI:** 10.64898/2026.09.27.754763

**Authors:** Fabrizio Musacchio, Martin Fuhrmann

## Abstract

Competitive spiking networks can form sensory representations through local synaptic plasticity and lateral inhibition. A canonical example is the Diehl–Cook architecture, in which Poisson-encoded sensory inputs drive excitatory neurons through spike-timing-dependent plasticity (STDP), while a separate inhibitory population imposes competition between excitatory units. In most implementations this competitive circuit is fixed, although inhibitory synapses are plastic in cortical circuits and contribute to excitation–inhibition balance. It therefore remains unclear under which conditions biologically motivated inhibitory plasticity can replace fixed lateral competition without disrupting representation learning. Here we study this question in a competitive STDP network trained on MNIST. We compare fixed inhibition with a Vogels-style inhibitory STDP rule, slow homeostatic inhibition, and a budget-constrained slow homeostatic rule that conserves the total inhibitory input received by each excitatory neuron. A fixed-inhibition operating-point map showed that stability depends on the dynamical regime: 56% of tested fixed-inhibition runs completed training, whereas stronger inhibition at higher input gain produced runaway activity. The reference regime *g* = 2.0, *w*^*IE*^ = 10.0, selected for subsequent comparisons, combined complete stability across replicate initializations with high classification performance (84.3±0.6% after 30,000 training examples). In this regime, the budget-constrained rule was stable in all replicate runs when its inhibitory budget matched the fixed-inhibition reference (*B*_*I*_ = 3990), reaching 83.5 ± 0.6% accuracy with firing-rate and inhibitory-conductance statistics close to fixed inhibition, whereas a mismatched budget (*B*_*I*_ = 7980) produced runaway activity in every replicate. Parameter maps showed that Vogels-derived iSTDP was broadly but not uniformly stable (90% runs completed), slow homeostatic inhibition was even more broadly stable (96%). Budget-constrained stability depended strongly on the conserved budget, i.e., low and matched budgets were stable or nearly stable, intermediate budgets were mixed, and *B*_*I*_ = 7980 failed in all tested runs. These results show that inhibitory plasticity in competitive STDP networks is not governed by a local homeostatic update alone. Stable plastic competition depends jointly on the operating regime and on an inhibitory budget that fixes the total strength of competition while allowing local inhibitory weights to adapt. Mechanistically, budget matching separates local redistribution of inhibitory weights from changes in total inhibitory drive, preventing plasticity from shifting the network away from a stable competitive state. This identifies inhibitory budget matching as a practical design principle for replacing fixed inhibition with plastic, homeostatically regulated competition in STDP-based spiking networks.

## Introduction

Spike-timing-dependent plasticity (STDP) provides a local learning rule by which synapses can strengthen or weaken as a function of relative pre- and postsynaptic spike timing [1, 2, 3]. In excitatory networks, such local Hebbian rules naturally promote competition, i.e., neurons that respond early and consistently to a pattern strengthen their afferent synapses and become more likely to win future competitions [2]. This principle has made STDP an attractive mechanism for unsupervised representation learning in spiking neural networks (SNN), from early visual-feature learning models to probabilistic cortical microcircuits and benchmark sensory classifiers [4, 5, 6, 7, 8]. The Diehl–Cook network is a particularly influential example of this approach [7]. It combines Poisson-encoded sensory input, plastic input-to-excitatory synapses, adaptive excitatory thresholds, and a separate inhibitory population that implements lateral competition. When evaluated on MNIST, the architecture develops class-selective excitatory responses without supervised weight updates, and classification can be performed post hoc by assigning each excitatory neuron to the class for which it fires most strongly. This benchmark use is important, but the architectural principle is more general than digit recognition: local feedforward STDP forms stimulus-selective excitatory assemblies, while inhibition shapes the competition between them. Related spiking sparse-coding models show that separate inhibitory populations can actively decorrelate excitatory units and support sparse receptive-field formation, rather than merely suppressing activity [9].

A crucial design choice in this architecture is that the recurrent competitive circuit is fixed, i.e., excitatory-to-inhibitory and inhibitory-to-excitatory connections establish the winner-take-all-like interaction, while the main plasticity occurs in the feedforward input synapses. This fixed-inhibition assumption is computationally convenient but biologically incomplete. Inhibitory synapses in cortical circuits are plastic, and inhibitory plasticity is widely thought to contribute to excitation–inhibition balance, receptive-field refinement, and stable activity regulation [10, 11, 12, 13, 14, 15, 16, 17]. Balanced excitatory and inhibitory interactions are also central to theoretical descriptions of cortical network dynamics [18, 19, 20]. In recurrent spiking models, inhibitory plasticity can stabilize firing rates and memory dynamics by adjusting inhibition in response to local activity [12, 15, 21]. Interacting excitatory and inhibitory plasticity can also shape receptive-field formation, stimulus-specific competition, and inhibitory assembly structure [22, 23, 24, 25]. More broadly, stable learning in spiking networks is rarely produced by a single plasticity mechanism. Hebbian plasticity is powerful precisely because it amplifies correlations, but this also makes it prone to runaway unless it is coupled to normalization, synaptic scaling, or other homeostatic constraints [26, 27, 28, 29, 17, 30]. Recent theory has sharpened this point by showing that functional recurrent excitatory–inhibitory networks can self-organize when Hebbian changes at multiple synapse types compete for limited synaptic resources [31]. These observations motivate a simple but underexplored question: can a biologically plausible inhibitory learning rule replace fixed lateral inhibition in a competitive STDP-based learning architecture? The answer is not obvious. The inhibitory STDP rule introduced by Vogels et al. [12] was developed to balance excitation and inhibition in recurrent networks, but a Diehl–Cook-style architecture already contains adaptive thresholds, feedforward weight normalization, strong lateral competition, and stimulus-driven winner selection. Previous theory has shown that the equilibria of Hebbian and STDP systems depend strongly on the constraints under which weights are allowed to evolve [32, 33, 34, 31]. A local inhibitory rule that is stabilizing in one dynamical context may therefore become destabilizing in another.

Here we examine how the dynamical regime of a competitive STDP network shapes the stability of inhibitory plasticity. We first ask which fixed-inhibition regimes are stable across replicate initializations. We then compare inhibitory learning rules within a stable baseline regime, separating three hypotheses. First, local inhibitory STDP adapted from Vogels et al. [12] may be sensitive to the competitive regime. Second, slow homeostatic inhibitory learning may avoid spike-by-spike instabilities but can still drift if its total inhibitory strength is unconstrained. Third, an explicit inhibitory budget constraint may stabilize plastic inhibition, provided that the budget matches a stable fixed-inhibition operating point. We therefore do not ask only whether inhibition can be made plastic, but also what must be conserved for plastic inhibition to remain compatible with stable competition. In this view, the key control variable is not a single local update term in isolation, but the interaction between local inhibitory adaptation and the overall strength of the competitive circuit.

## Methods

### Network architecture

We studied a competitive spiking network with an input population *X*, an excitatory population *E*, and an inhibitory population *I* (Fig. 1a). We used MNIST as a standard benchmark for unsupervised sensory representation learning and comparability with the Diehl–Cook model [6, 7]. Each MNIST image is a 28 × 28 grayscale digit image. Images were flattened into *N*_*X*_ = 784 input channels and converted into independent Poisson spike trains, so the network received a rate-coded input vector rather than an explicitly two-dimensional convolutional representation. For pixel *k* with intensity *p*_*k*_ ∈ [0, 255], the input rate was

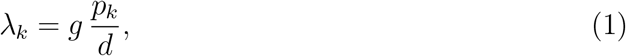

where *g* is a scalar input gain and *d* is a fixed intensity divisor. The input-to-excitatory projection *W*^*XE*^ was plastic. The excitatory-to-inhibitory projection *W*^*EI*^ was one-to-one and fixed. The inhibitory-to-excitatory projection *W*^*IE*^ was either fixed or plastic depending on the experimental condition. Self-inhibition was excluded by setting 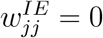 when inhibitory and excitatory populations had matched size.

**Figure 1:**
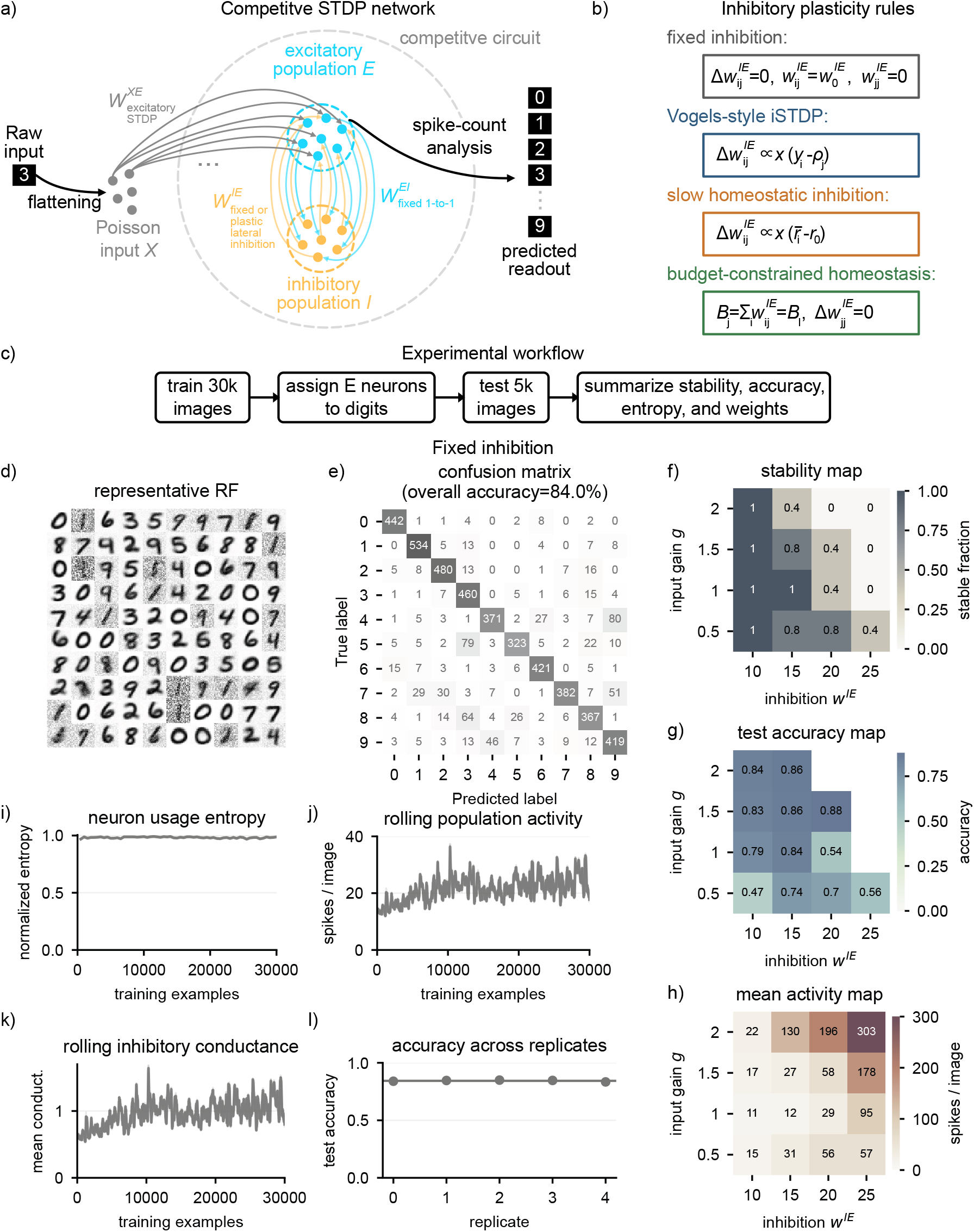
Competitive STDP network, experimental workflow, and fixed-inhibition operating regime. (a) MNIST images are flattened into Poisson input neurons, which project to an excitatory population through plastic excitatory STDP. Excitatory neurons recruit a matched inhibitory population that sends lateral inhibitory input back to the excitatory population. The central question is which inhibitory plasticity rule can preserve this competitive circuit without runaway activity, collapse, or weight saturation. (b) We compared fixed inhibition, inhibitory STDP adapted from Vogels et al. [12], slow homeostatic inhibition, and budget-constrained slow homeostatic inhibition. (c) Each run followed the same 30,000-example training, label-assignment, 5000-example testing, and summary workflow. (d) Representative input-to-excitatory receptive fields from the selected stable fixed-inhibition run. (e) Representative post hoc digit-readout confusion matrix from the same run. (f) Fraction of stable fixed-inhibition replicates across input gain and inhibitory strength. (g) Mean test accuracy among non-aborted fixed-inhibition runs. (h) Mean excitatory spikes per image across the same fixed-inhibition sweep. (i) Neuron usage entropy over training for the five replicates at the selected stable fixed-inhibition operating point. (j) Rolling mean excitatory spikes per image at the selected operating point. (k) Rolling mean inhibitory conductance at the selected operating point. (l) Test accuracy across replicate initializations at the selected stable operating point *g* = 2.0, *w*^*IE*^ = 10.0.

Excitatory and inhibitory neurons were conductance-based leaky integrate-and-fire units. For an excitatory neuron *j*, membrane potential evolved as

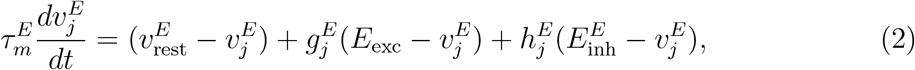

where 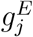 and 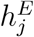 are excitatory and inhibitory conductances. When 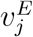 crossed its adaptive threshold, the neuron emitted a spike, entered a refractory state, and its adaptive threshold component *θ*_*j*_ increased by Δ*θ*. The effective threshold was

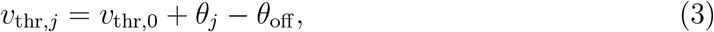

where the adaptive component followed

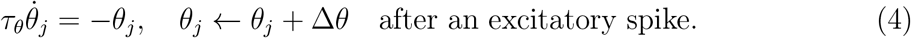

Inhibitory neurons followed the same conductance-based structure without adaptive thresholds. All state variables were advanced with a forward Euler scheme with time step Δ*t* = 0.5 ms. Each stimulus was presented for 350 ms followed by a 150 ms resting period. Conductances and spike traces were decayed exponentially at each time step, and threshold adaptation was integrated with the same Euler clock.

### Excitatory STDP and label assignment

The input-to-excitatory synapses followed the Diehl–Cook-style excitatory STDP mechanism [7]. For input neuron *k* and excitatory neuron *j*, presynaptic traces *a*_*k*_ and postsynaptic traces *b*_*j*_, *c*_*j*_ decayed exponentially and were incremented by spikes,

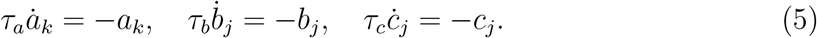

At input spikes, the synapse was depressed in proportion to the postsynaptic trace,

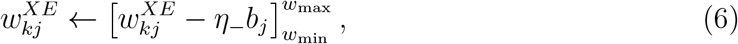

whereas at excitatory spikes it was potentiated in proportion to the presynaptic trace and the postsynaptic spike trace immediately before reset,

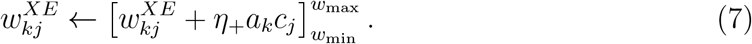

After stimulus presentations, incoming input weights were normalized so that each excitatory neuron received a comparable total feedforward drive,

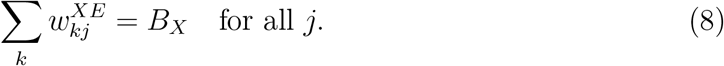

This normalization is important because otherwise a small number of neurons can dominate the input representation.

Classification was evaluated after unsupervised training by assigning each excitatory neuron to the digit class that elicited its highest average spike count on the training set. For each test image, class evidence was computed from the spike counts of neurons assigned to that class. The predicted digit was the class with maximal evidence.

### Inhibitory plasticity rules

We represented inhibitory weights as non-negative magnitudes 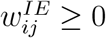 from inhibitory neuron *i* to excitatory neuron *j*. These weights did not add a separate state variable to the membrane equation. Instead, they scaled the inhibitory conductance increments delivered to excitatory neurons when inhibitory spikes occurred. After a plastic update changed 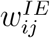, subsequent inhibitory spikes therefore entered the leaky integrate-and-fire dynamics through the inhibitory conductance term 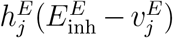. For spike-based inhibitory STDP, inhibitory presynaptic traces *x*_*i*_ and excitatory postsynaptic traces *y*_*j*_ obeyed

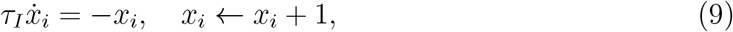

after inhibitory spikes and

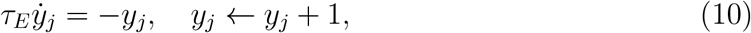

after excitatory spikes.

The inhibitory STDP rule adapted from Vogels et al. [12] applied a depressive offset at inhibitory presynaptic spikes and potentiated inhibition when pre- and postsynaptic activity were coincident. In event-based form,

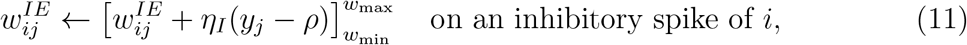

and

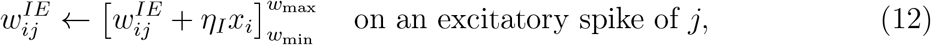

where 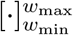 denotes clipping.

We also tested a slow homeostatic inhibitory rule. For stimulus *s*, let 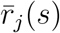 denote the stimulus-evoked spike count of excitatory neuron *j* expressed as a stimulus-rate summary, and let 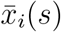 denote the integrated inhibitory presynaptic activity over the same stimulus window. The slow rule updated

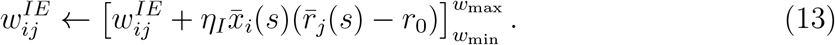

This rule removes fast coincidence potentiation and instead adjusts inhibition from stimulus-scale deviations around a target rate *r*_0_.

Finally, the budget-constrained slow homeostatic rule applied the same local update but followed it with column-wise normalization (Fig. 1b):

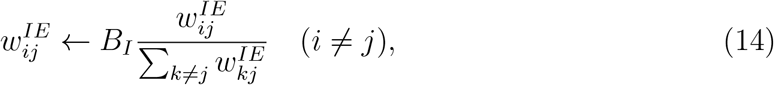

with 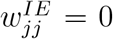. The scalar *B*_*I*_ is the inhibitory budget received by each excitatory neuron. In the subsequent experiments, we distinguish a mismatched budget *B*_*I*_ = 7980, corresponding to an average inhibitory weight near 20, from a matched budget *B*_*I*_ = 3990, corresponding to the stable fixed-inhibition operating point with average inhibitory weight near 10 (Fig. S2).

### Budget constraint as a competition-control principle

Normalization and conservation constraints are a classical way to make Hebbian learning stable without removing its local correlation-sensitive component [32, 33]. They also determine the equilibria reached by temporally asymmetric Hebbian plasticity and STDP-like rules [34]. In biological networks, related stabilizing principles appear as homeostatic synaptic scaling and other cell-wide regulatory processes that compensate for correlation-amplifying Hebbian changes [26, 27, 17]. Recent work on recurrent excitatory–inhibitory networks further showed that synapse-type-specific competition for limited synaptic resources can stabilize simultaneous Hebbian plasticity at multiple synapse classes and support functional receptive-field organization [31]. Our inhibitory budget normalization should be read in this tradition, but with a narrower purpose. Rather than modeling the development of a fully plastic recurrent circuit, it tests whether a fixed inhibitory-to-excitatory projection in a competitive spiking STDP architecture can be replaced by local inhibitory plasticity without changing the total competitive drive. Thus, the budget is not only a numerical bound on inhibitory weights, but a constraint on the total inhibition received by each excitatory neuron.

In a competitive excitatory–inhibitory circuit, the column sum

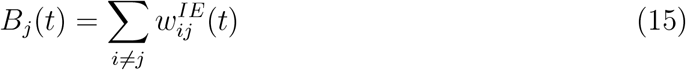

sets the total inhibitory drive available to suppress excitatory neuron *j* after the inhibitory population has been recruited. A purely local inhibitory update changes both the relative distribution of inhibitory weights and the total amount of inhibition received by a postsynaptic neuron,

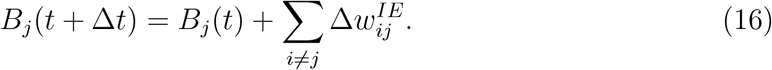

Thus, local homeostatic plasticity can unintentionally move the network between competitive regimes. Changing *B*_*j*_ changes how strongly excitatory neurons recruit lateral suppression and how much input drive, threshold adaptation, and recurrent excitation– inhibition feedback are needed to elicit responses. Too little total inhibition can weaken competition, whereas too much total inhibition can push the network toward high-gain response regimes that end in runaway or collapse. In a winner-take-all-like STDP network, the relevant control problem is therefore not only whether each synapse follows a stabilizing local rule, but whether the population remains near a viable total-inhibition regime. This differs from unconstrained clipping. Clipping limits individual weights, whereas the budget fixes the aggregate inhibitory drive while still allowing local weights to redistribute.

The budget-constrained rule separates these two degrees of freedom. The local slow-homeostatic update redistributes inhibition across inhibitory inputs according to activity, while the normalization step enforces *B*_*j*_ = *B*_*I*_ for every excitatory neuron. If *B*_*I*_ is chosen from a stable fixed-inhibition operating point, inhibitory plasticity can alter the pattern of inhibitory weights without changing the total competition strength. If *B*_*I*_ is mismatched, the same normalization can stabilize the wrong regime. This distinction motivated the matched- and mismatched-budget comparisons in the results.

### Stability and representation metrics

The main stability metric was whether a run completed the prescribed training subset without triggering an early-abort criterion. A run was labeled as runaway if the excitatory population emitted more than 5000 spikes per image for 20 consecutive examples. Runs were also aborted if the mean adaptive threshold exceeded 200, if more than 20% of inhibitory synapses reached the upper weight bound, or if silent-collapse retries occurred for 100 consecutive examples. In retry cases, the dynamic state and plastic weights were restored before the same stimulus was presented again at higher input intensity, so failed low-spike presentations did not cumulatively update the network before the retry.

For each condition we summarized mean excitatory spikes per image, mean inhibitory conductance, the mean adaptive threshold, and statistics of *W*^*IE*^. Let *n*_*j*_(*s*) denote the number of spikes emitted by excitatory neuron *j* during stimulus presentation *s*. Population activity was summarized as the excitatory spike count per image,

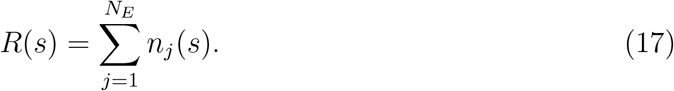

We used two related time-course summaries. Rolling traces were computed directly from single-stimulus recordings. Rolling excitatory spike counts are moving averages of *R*(*s*) over 100 consecutive training examples, and rolling inhibitory conductance was computed analogously from the mean inhibitory conductance recorded for each stimulus. Windowed summaries were computed from the training-window metrics saved during the run, using the same non-overlapping windows that were used for condition-level monitoring. Thus, windowed panels show coarser training-window estimates of usage entropy, population activity, or inhibitory conductance, whereas rolling panels show smoothed single-example trajectories. Both summaries were used only for visualization. Condition-level statistics were computed from the recorded training-window metrics and final run summaries.

We quantified neuron usage by the entropy of the excitatory spike distribution. For an analysis window S, let

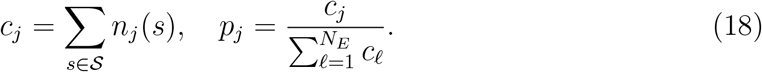

Normalized neuron-usage entropy was then

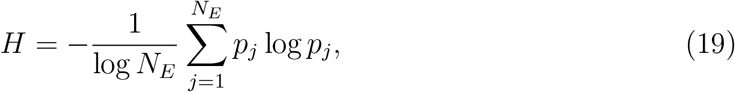

where terms with *p*_*j*_ = 0 were omitted. Thus, *H* = 1 indicates that spikes were distributed uniformly across excitatory neurons, while lower values indicate concentration of activity in a smaller subset of neurons. We also measured the firing-rate coefficient of variation across excitatory neurons. A neuron was counted as dead within a window if *c*_*j*_ = 0, and the dead-neuron fraction was the fraction of excitatory neurons satisfying this criterion. Unless stated otherwise, final usage, dead-neuron, conductance, and threshold summaries refer to the final training window or final network state of each run and are reported as mean ± standard deviation across replicate initializations.

We use the term *map* for systematic parameter-grid experiments in which two control parameters are varied and each grid point is summarized across replicate initializations. For fixed inhibition, the map varied input gain *g* and inhibitory strength *w*^*IE*^ and summarized stable-replicate fraction, test accuracy, and mean population activity. For the Vogels-derived inhibitory STDP rule, the map varied the inhibitory learning rate *η*_*I*_ and target offset *ρ* and summarized stable-replicate fraction, number of training examples processed before completion or abort, and replicate-to-replicate variability. In these maps, each colored cell therefore represents a regime-level summary rather than a single trajectory.

To compare the internal organization of learned weights, we also computed pairwise Pearson correlations between excitatory-neuron weight columns. For *W*^*XE*^, each column is the feedforward input profile of one excitatory neuron and therefore summarizes its receptive-field similarity to other excitatory neurons. For *W*^*IE*^, each column is the inhibitory input profile received by one excitatory neuron. Because self-inhibition is excluded, the diagonal entries of square *W*^*IE*^ matrices were replaced by the corresponding column mean before computing correlations, so that structural zeros did not dominate the inhibitory-profile comparison. Heatmaps show these correlation matrices after sorting excitatory neurons by their post hoc digit assignment. We further summarized the median inhibitory-profile correlation within the same assigned digit and between different assigned digits. Finally, we quantified feedforward–inhibitory coupling by correlating the upper-triangular entries of the *W*^*XE*^ similarity matrix with the corresponding entries of the *W*^*IE*^ similarity matrix. These weight-correlation diagnostics were used to test whether successful plastic-inhibition runs merely reproduced output-level behavior or also reorganized the hidden inhibitory connectivity.

### Experimental design

All primary experiments used *N*_*E*_ = *N*_*I*_ = 400 excitatory and inhibitory neurons, one epoch of 30,000 MNIST training examples, 5000 test examples, and five random initializations. MNIST contains 60,000 training images and 10,000 test images [6]. We used fixed half-size train and test subsets to make replicate regime sweeps and inhibitory-plasticity comparisons computationally tractable while retaining substantially more training than the initial proof-of-concept runs. A separate long fixed-inhibition run with 60,000 training examples and two epochs showed stable activity statistics, broad neuron usage, and little improvement in test accuracy after the first full epoch (Fig. S1). We therefore used the 30,000-example protocol as a compromise between numerical cost and stable representation formation. Each run followed the same training, post hoc label-assignment, testing, and summary workflow (Fig. 1c). The goal of these experiments was to characterize stability mechanisms under matched conditions rather than to optimize final MNIST benchmark performance. Accordingly, a plastic inhibitory rule was considered successful if it preserved the fixed-reference operating regime, i.e., comparable activity, representation, and readout statistics, while replacing fixed *W*^*IE*^ with adaptive inhibition. We first performed a fixed-inhibition operating-regime sweep over input gain *g* ∈ {0.5, 1.0, 1.5, 2.0} and initial inhibitory strength *w*^*IE*^ ∈ {10, 15, 20, 25}. We then compared inhibitory plasticity rules in the stable fixed-inhibition reference regime *g* = 2.0, *w*^*IE*^ = 10.0. The two Vogels-derived conditions used in this direct comparison, *ρ* = 0.01 and *ρ* = 0.03, probed sensitivity to the target-offset term at fixed learning rate and weight bound. Additional stability maps tested the Vogels-derived, slow-homeostatic, and budget-constrained rules over their rule-specific control parameters. The main simulation and comparison parameters are summarized in Table 1.

**Table 1:** Primary simulation and comparison parameters. Values apply to all primary runs unless explicitly listed as swept or condition-specific. The readout is a post hoc class-evidence calculation from excitatory spike counts, not a trained output-neuron layer.

| Quantity | Value | Notes |
| --- | --- | --- |
| **Network and numerical settings** |  |  |
| Input population | $N_X = 784$ | Flattened $28 \times 28$ MNIST images encoded as independent Poisson spike trains. |
| Excitatory and inhibitory populations | $N_E = N_I = 400$ | Matched populations with excluded self-inhibition, $w_{jj}^{IE} = 0$. |
| Readout | 10 class evidence values | Post hoc label assignment from excitatory spike counts; no plastic output layer. |
| Numerical integration | $\Delta t = 0.5$ ms | Forward Euler integration for membrane variables, conductances, traces, and adaptive thresholds. |
| Stimulus protocol | 350 ms stimulus, 150 ms rest | One image presentation per training or test example. |
| Training and testing subsets | 30,000 train, 5000 test examples | One epoch, five replicate initializations; justified by the long fixed-inhibition diagnostic in Fig. S1. |
| Excitatory threshold adaptation | $\Delta\theta = 0.05$ mV, $\tau_\theta = 10^7$ ms | Adaptive threshold increment and slow decay. |
| Excitatory STDP traces | $\tau_a = 20$ ms, $\tau_b = 20$ ms, $\tau_c = 40$ ms | Input-to-excitatory STDP; incoming $W^{XE}$ columns normalized after stimulus presentations. |
| **Fixed competitive circuit and operating-regime sweep** |  |  |
| Fixed recurrent excitation | $w^{EI} = 10.4$ | One-to-one excitatory-to-inhibitory projection. |
| Fixed recurrent inhibition | $w^{IE} = 10, 15, 20, 25$ | Initial fixed-inhibition strengths tested in the operating-regime sweep. |
| Input gain sweep | $g \in \{0.5, 1.0, 1.5, 2.0\}$ | Five replicate initializations per parameter pair. |
| Reference operating point | $g = 2.0$, $w^{IE} = 10.0$ | Stable fixed-inhibition regime used for direct inhibitory-plasticity comparisons. |

**Table 1: Primary simulation and comparison parameters** (continued).
| Quantity | Value | Notes |
| --- | --- | --- |
| **Direct inhibitory-plasticity comparison** |  |  |
| Compared conditions | fixed; Vogels $\rho = 0.01$; Vogels $\rho = 0.03$; slow; budget-constrained | All compared in the selected fixed-inhibition reference regime with five replicate initializations. |
| Inhibitory spike traces | $\tau_I = \tau_E = 20$ ms | Used for the Vogels-derived inhibitory STDP rule. |
| Inhibitory learning rate and bounds | $\eta_I = 3 \cdot 10^{-4}$, $w_{\max}^{IE} = 25$ | Used for the direct inhibitory-plasticity comparison. |
| Slow homeostatic target | $r_0 = 0.1$ Hz | Stimulus-scale target for the direct slow-homeostatic comparison. |
| Budget-constrained inhibition | $B_I = 3990$ | Matched budget corresponding to $\bar{w}^{IE} \approx 10$. |
| Mismatched-budget control | $B_I = 7980$ | Negative-control budget corresponding to $\bar{w}^{IE} \approx 20$. |
| **Inhibitory-plasticity stability maps** |  |  |
| Vogels-derived map | $\eta_I \in \{10^{-5}, 3 \cdot 10^{-5}, 10^{-4}, 3 \cdot 10^{-4}, 10^{-3}\}$; $\rho \in \{0.01, 0.03, 0.1, 0.3\}$ | Five replicate initializations per parameter pair; $w_{\max}^{IE} = 25$. |
| Slow-homeostatic map | $\eta_I \in \{10^{-5}, 3 \cdot 10^{-5}, 10^{-4}, 3 \cdot 10^{-4}, 10^{-3}\}$; $r_0 \in \{0.03, 0.1, 0.3, 1.0\}$ Hz | Five replicate initializations per parameter pair. |
| Budget-constrained map | $\eta_I \in \{10^{-5}, 3 \cdot 10^{-5}, 10^{-4}, 3 \cdot 10^{-4}, 10^{-3}\}$; $B_I \in \{1995, 3990, 5985, 7980\}$ | Five replicate initializations per parameter pair. |
| **Retry and abort safeguards** |  |  |
| Retry handling | rollback enabled | Dynamic state and plastic weights were restored before increasing input intensity after a failed retry. |
| Rest-period inhibitory plasticity | disabled | iSTDP updates were not applied during the post-stimulus resting period. |
| Runaway criterion | $> 5000$ excitatory spikes for 20 consecutive examples | Detected sustained high-activity failures. |
| Silent-collapse criterion | 100 consecutive retry-cap examples | Retry cap was 25 attempts per image; rollback was applied before each retry. |
| Threshold and saturation criteria | $\bar{\theta} > 200$ or $> 20\%$ of inhibitory synapses at $w_{\max}^{IE}$ | Detected excessive threshold growth or inhibitory-weight saturation. |

## Results

### Fixed-inhibition operating-regime analysis

To interpret inhibitory plasticity against a controlled baseline, we first established a stable fixed-inhibition operating point for the competitive circuit, following the workflow illustrated in Fig. 1c and using the shared simulation parameters summarized in Table 1. In the selected fixed-inhibition run, the input-to-excitatory receptive fields contained many clear digit-like templates, interspersed with weaker or noisier fields that remained less specialized (Fig. 1d). This pattern is qualitatively consistent with Diehl–Cook-style unsupervised STDP networks, which form class-selective receptive fields but do not require every excitatory neuron to become an equally clean prototype [7]. The corresponding post hoc label-assignment readout produced a structured confusion matrix rather than random class evidence (Fig. 1e). Thus, the baseline entered the expected unsupervised representation-learning regime.

Quantitatively, the fixed reference reached 84.3 ± 0.6% test accuracy after 30,000 training examples. This is below the strongest Diehl–Cook benchmark configurations, which reported higher MNIST performance after extensive model-specific tuning and scaling, but it is in the same functional range as a working medium-sized STDP recognizer and is sufficient for the present stability comparisons [7]. Extending the baseline to the full 60,000 training examples for two epochs did not transform the qualitative regime into a substantially different benchmark solution. Here, test accuracy was 86.4% after the first epoch and 85.3% after the second, and feedforward weights remained strongly correlated between epochs (Fig. S1c,d). The remaining gap to the best reported Diehl–Cook-style results therefore more likely reflects implementation and optimization choices, including the Euler integration scheme, finite replicate-sweep protocol, reduced benchmark subset, and absence of task-specific hyperparameter optimization, than a failure of the baseline to learn useful representations. We therefore treated the residual noisy receptive fields and the reference-regime accuracy as a sufficient, stable sanity benchmark for comparing inhibitory plasticity rules.

We then mapped the fixed-inhibition circuit before interpreting any plastic inhibitory rule. The regime maps in Fig. 1f–h summarize all fixed-inhibition grid runs. In each map, the horizontal axis is the fixed inhibitory strength *w*^*IE*^, the vertical axis is the input gain *g*, and color encodes the aggregate outcome at each parameter pair. Darker cells in the stability map indicate that more replicate initializations completed training, while pale or white cells indicate unstable regimes in which no replicate completed training. Across 80 fixed-inhibition runs, stability depended jointly on input gain and inhibitory strength (Fig. 1f). Overall, 45*/*80 runs completed training. The zero-stability cells at high inhibition and higher input gain show that fixed inhibition is not automatically safe, i.e., the same competitive architecture can enter runaway regimes when the input drive and inhibitory strength are mismatched.

The corresponding accuracy and activity maps in Fig. 1g,h show why the stable region cannot be reduced to a trivial low-activity regime. Several low- and intermediate-inhibition conditions remained both stable and accurate, while stronger inhibition at higher input gain was associated with high activity among completed or aborted runs. We selected *g* = 2.0 and *w*^*IE*^ = 10.0 as the reference operating point because it completed all 30,000 training examples in all five replicate initializations, yielded 84.3 ± 0.6% test accuracy, and maintained a mean activity of 21.51 ± 0.05 excitatory spikes per image (Fig. 1l).

The regime maps identify where stable fixed-inhibition learning is possible, but they do not show whether the selected point remains stable throughout training. We therefore examined the five replicate trajectories for the selected operating point only (Fig. 1i–k). Neuron usage remained broad over training (*H* = 0.9977 ± 0.0001 at the end of training; Fig. 1i), rolling excitatory spike counts remained in the same activity band used to select the reference regime (Fig. 1j), and inhibitory conductance showed no progressive drift (Fig. 1k). The final dead-neuron fraction was 0.0 ± 0.0 across the five replicate initializations. The final adaptive threshold, averaged first over the 400 excitatory neurons within each replicate and then summarized across replicates, was 32.08±0.04. We therefore used the validated reference operating point (*g* = 2.0, *w*^*IE*^ = 10.0) as the baseline for subsequent inhibitory-plasticity comparisons.

### Vogels-derived inhibitory STDP in the reference regime

We next asked whether a spike-based inhibitory STDP rule adapted from Vogels et al. [12] could replace fixed inhibition while preserving the same competitive learning workflow. The rule was evaluated in the fixed-inhibition reference regime using the same training, label-assignment, testing, and summary procedure as the baseline (Fig. 1c; Table 1), but with *W*^*IE*^ updated by the event-based inhibitory-spike and excitatory-spike updates in Eqs. (11) and (12). We tested two values of the homeostatic target offset in the inhibitory presynaptic update, *ρ* = 0.01 and *ρ* = 0.03, to determine whether modest changes in this depressive offset affected stability when the learning rate and weight bound were held fixed. The representative diagnostics in Fig. 2 illustrate both the stable and unstable outcomes observed under this rule. The failed low-*ρ* example already showed disrupted receptive fields at abort (Fig. 2a), while completed low- and high-offset runs produced recognizable receptive fields and structured post hoc readout matrices (Fig. 2b–e). Their window-based entropy, population activity, and inhibitory-conductance traces remained in the stable reference regime, whereas the failed low-*ρ* trajectory revealed the delayed runaway mode (Fig. 2f,g,i,j,l).

**Figure 2:**
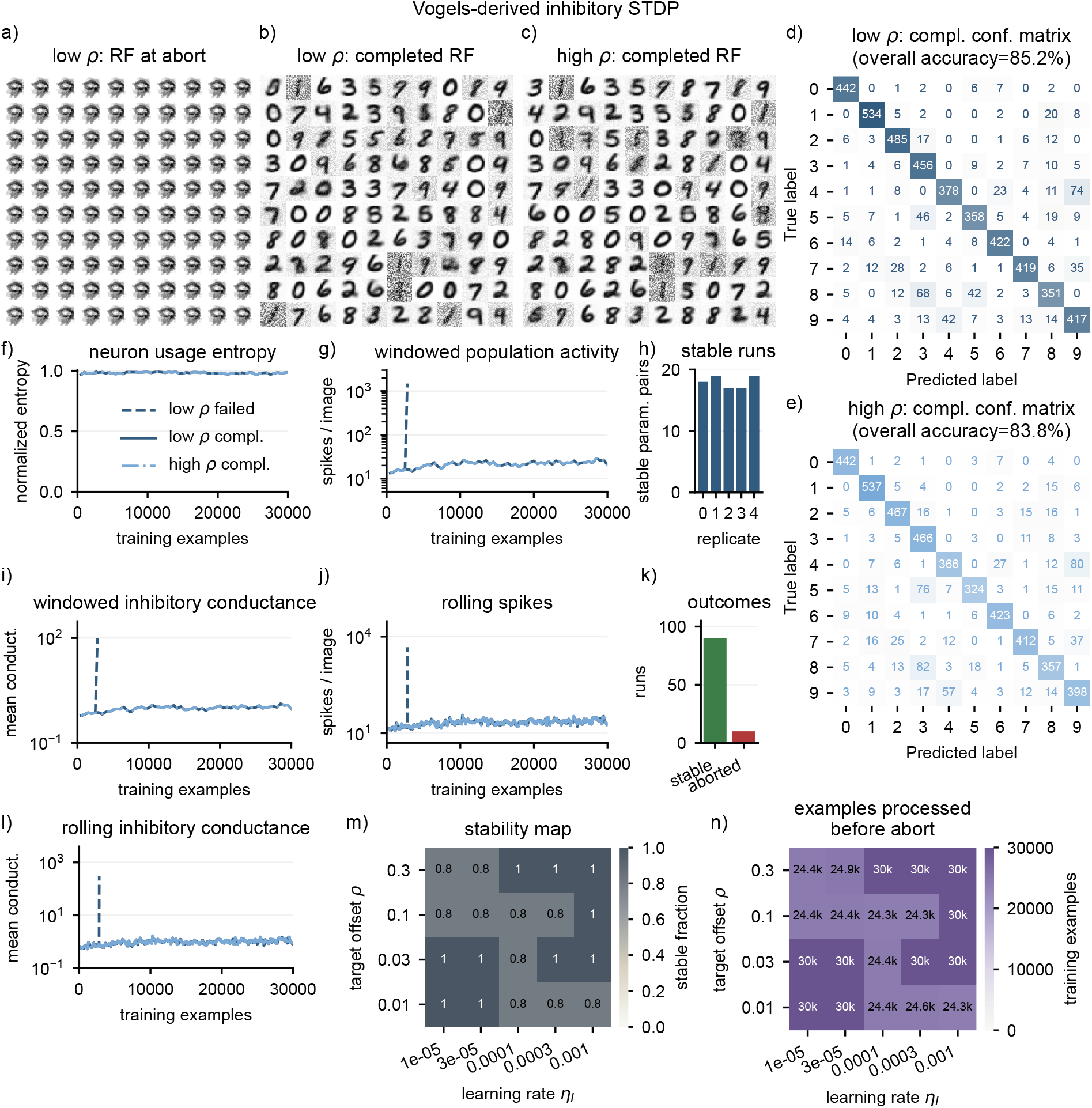
Vogels-derived inhibitory STDP can preserve the reference regime but remains parameter- and initialization-sensitive. (a) Receptive fields from a low-*ρ* run that aborted before testing. (b,c) Receptive fields from completed low-*ρ* and high-*ρ* runs. (d,e) Post hoc readout confusion matrices from the same completed low-*ρ* and high-*ρ* runs. No high-*ρ* abort panel is shown because all five direct-comparison high-*ρ* replicate initializations completed training. (f,g,i) Window-based neuron-usage entropy, excitatory population activity, and inhibitory conductance for the failed low-*ρ*, completed low-*ρ*, and completed high-*ρ* examples. (h) Number of stable parameter pairs per replicate initialization. (j,l) Rolling single-example spike-count and inhibitory-conductance trajectories for the same examples. (k) Total stable and aborted outcomes across the 100-run map. (m) Fraction of stable replicate runs across learning rate *η*_*I*_ and target offset *ρ*. (n) Mean number of training examples processed before completion or abort.

In the summary comparison in Fig. 3 we show how these runs related to the fixed reference and the other inhibitory-plasticity rules. The rule-stability summary in Fig. 3a shows that the lower-offset condition (*ρ* = 0.01) completed four of five replicate initializations, while the higher-offset condition (*ρ* = 0.03) completed all five. Classification performance among completed runs remained close to fixed inhibition for both offsets (Fig. 3c). This was not achieved by shifting the network into a different firing-rate regime, i.e., mean activity for both offsets remained near 21.5 spikes per image (Fig. 3d). Inhibitory conductance trajectories nearly overlapped the fixed-inhibition trajectory throughout training (Fig. 3e), and final mean inhibitory weights remained close to the reference inhibitory strength (Fig. 3f). Thus, in the matched reference regime, Vogels-derived inhibitory STDP preserved the same functional operating point, consistent with the goal of replacing fixed inhibition without shifting the network into a qualitatively different activity state.

**Figure 3:**
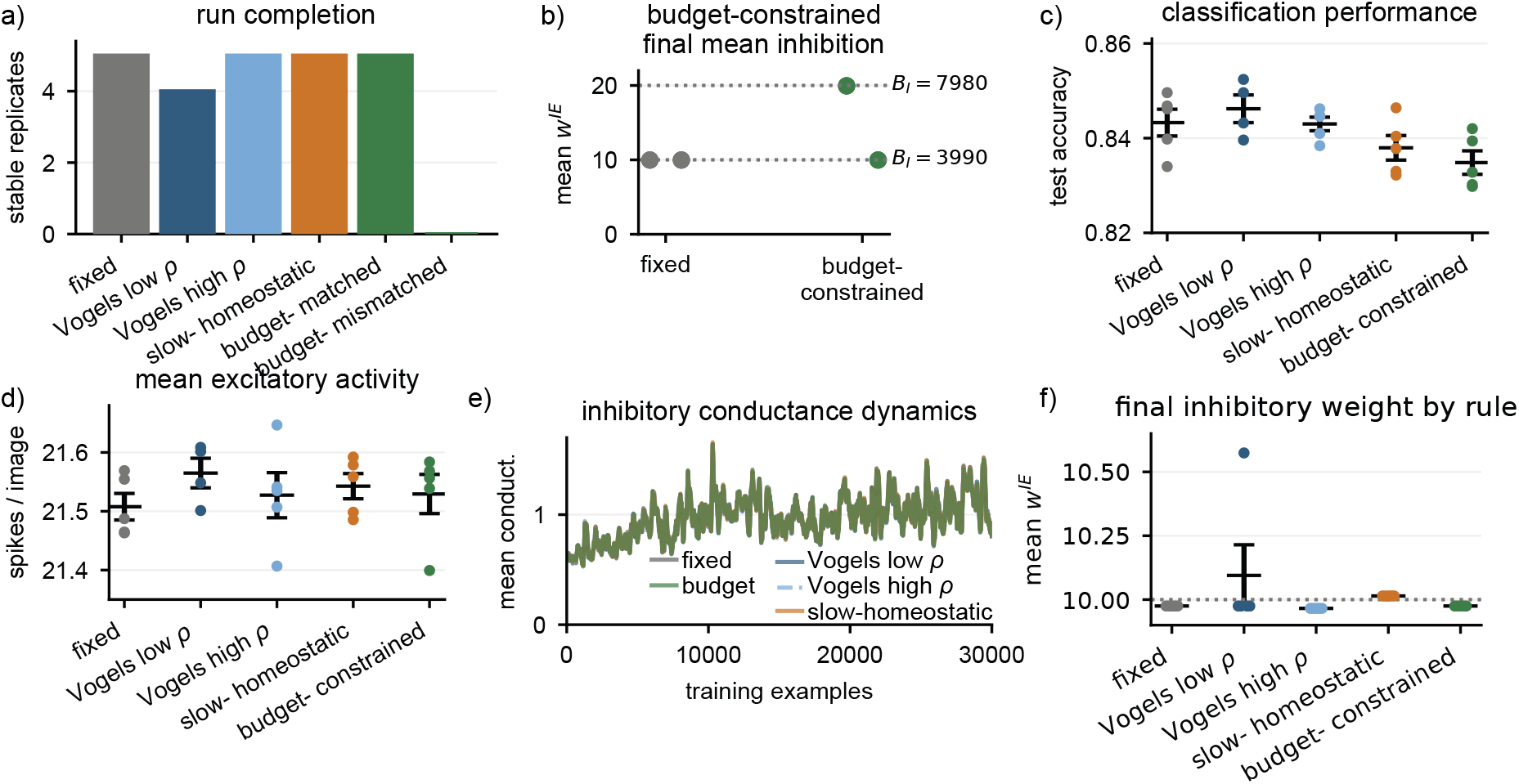
Inhibitory plasticity depends on the conserved inhibitory budget. (a) Stable replicate runs for fixed inhibition and inhibitory plasticity rules in the matched-budget/reference regime, plus the negative control in which the budget-constrained slow-homeostatic rule used a mismatched inhibitory budget (*B*_*I*_ = 7980). (b) Final mean inhibitory weight for fixed and budget-constrained inhibition. The mismatched budget-constrained rule enforces an inhibitory budget corresponding to mean inhibition near 20, while the matched rule preserves mean inhibition near 10. (c) Test accuracy among completed runs in the matched-budget regime. (d) Mean excitatory spikes per image among completed runs in the matched-budget regime. (e) Inhibitory conductance dynamics during training for fixed inhibition, inhibitory STDP adapted from Vogels et al. [12], unconstrained slow homeostasis, and budget-constrained slow homeostasis. (f) Final mean inhibitory weights for the same matched-budget method comparison.

This matched-reference comparison shows that Vogels-derived iSTDP can preserve the selected regime. It does not, however, show how stability is distributed across the rule’s own parameter space. We therefore mapped the rule across five learning rates, four target offsets, and five replicate initializations, using the parameter grid summarized in Table 1. In Fig. 2m, the horizontal axis is the inhibitory learning rate *η*_*I*_, the vertical axis is the target offset *ρ*, and color encodes the fraction of stable replicate initializations at each parameter pair. Across the resulting 100 runs, 90 completed training, as summarized by the stable-versus-aborted outcome counts in Fig. 2k. Thus, Vogels-derived iSTDP was broadly compatible with the selected competitive regime. However, the ten aborted runs indicate that this compatibility was conditional rather than uniform, with stability still depending on parameter choice and replicate initialization.

The aborted runs were nevertheless informative. Six showed runaway failures and four showed silent-collapse failures detected by repeated retry-cap events. Because each replicate initialization was tested across 20 parameter pairs, the replicate summary in Fig. 2h shows that every initialization produced at least one failure, while no initialization accounted for all failures. Together with the stability map in Fig. 2m, this indicates that failures were distributed across both replicate initializations and parameter pairs rather than being confined to one bad seed or one simple monotonic corner of the grid. The processed-example heatmap in Fig. 2n shows that unstable conditions often ran for thousands of examples before failing, indicating delayed dynamical failure rather than immediate numerical incompatibility. A representative failed low-offset run illustrates this delayed runaway: activity remained near the stable band initially, then rapidly increased by orders of magnitude together with inhibitory conductance (Fig. 2a,f,g,i,j,l; Fig. 5a,b). These results indeed extend the baseline analysis by showing that local spike-based inhibitory plasticity can be compatible with competitive STDP, but only within an appropriate operating regime and parameterization.

### Slow homeostatic inhibitory plasticity in the reference regime

We then removed fast spike-coincidence potentiation and tested the stimulus-scale slow homeostatic update defined in Eq. (13). In the selected fixed-inhibition reference regime (*g* = 2.0, *w*^*IE*^ = 10.0), unconstrained slow homeostasis completed all five replicate initializations (Fig. 3a). Its test accuracy remained close to fixed inhibition (83.8 ± 0.6%; Fig. 3c), and its mean excitatory activity matched the narrow activity band shared by the other stable matched-budget conditions (21.54 ± 0.05 spikes per image; Fig. 3d). The inhibitory conductance trajectory in Fig. 3e closely overlapped the fixed and Vogels-derived trajectories throughout training, indicating that the slower update did not introduce a separate activity timescale or progressive conductance drift.

The main difference between slow homeostasis and fixed inhibition appeared in the inhibitory weights. In Fig. 3f, the final mean inhibitory weight stayed near the fixed reference (10.01 ± 0.0003), but this scalar summary hides the fact that the slow rule redistributed individual inhibitory synapses. The individual-weight and column-sum diagnostics in Fig. 5c,d separate these two levels of organization. Slow homeostasis and matched budget-constrained homeostasis both redistributed local inhibitory weights relative to the almost uniform fixed and Vogels-derived cases, but only the budget-constrained rule conserved the column-wise inhibitory budget. The weight-correlation diagnostics in Fig. 5e–l add a second view of the same mechanism. Feedforward weight correlations were nearly indistinguishable across methods (Fig. 5e), consistent with the similar receptive fields and readout performance. In contrast, inhibitory input-profile correlations separated fixed from plastic inhibition. Fixed inhibition produced nearly identical inhibitory profiles, whereas all plastic rules redistributed *W*^*IE*^ into non-uniform, label-structured profiles (Fig. 5f–k; Fig. S3). Within the same post hoc digit assignment, inhibitory profiles were positively correlated, while between different digit assignments they were weakly anticorrelated or near zero (Fig. 5k). Thus, negative *W*^*IE*^ profile correlations do not imply negative inhibition. They indicate complementary inhibitory fingerprints across excitatory neurons. The coupling between feedforward and inhibitory similarity further showed that excitatory neurons with similar *W*^*XE*^ profiles tended to receive more similar inhibitory profiles (Fig. 5l). The rule-specific diagnostics in Fig. 4 show that this hidden inhibitory reorganization was compatible with representation learning, since the completed run produced digit-like receptive fields and a structured confusion matrix (Fig. 4b,c). Its entropy, activity, and conductance traces also remained stable over training (Fig. 4d–h). The same figure also shows a separate early saturation example in which the slow-homeostatic update was pushed into inhibitory weight saturation rather than stable redistribution, with degraded receptive fields at abort and divergent activity/conductance dynamics (Fig. 4a,d–h). The stability map in Fig. 4i extends this direct comparison across *η*_*I*_ and *r*_0_: 96*/*100 slow-homeostatic runs completed training, and the four failures were isolated single-replicate failures rather than a contiguous unstable parameter region. The processed-example map in Fig. 4j shows that these failures occurred late in training, after roughly 24,000–25,000 examples on average at the affected grid points. Thus, slow homeostatic inhibition was sufficient to preserve stable competition across most of the tested parameter range, even without an explicit budget constraint.

**Figure 4:**
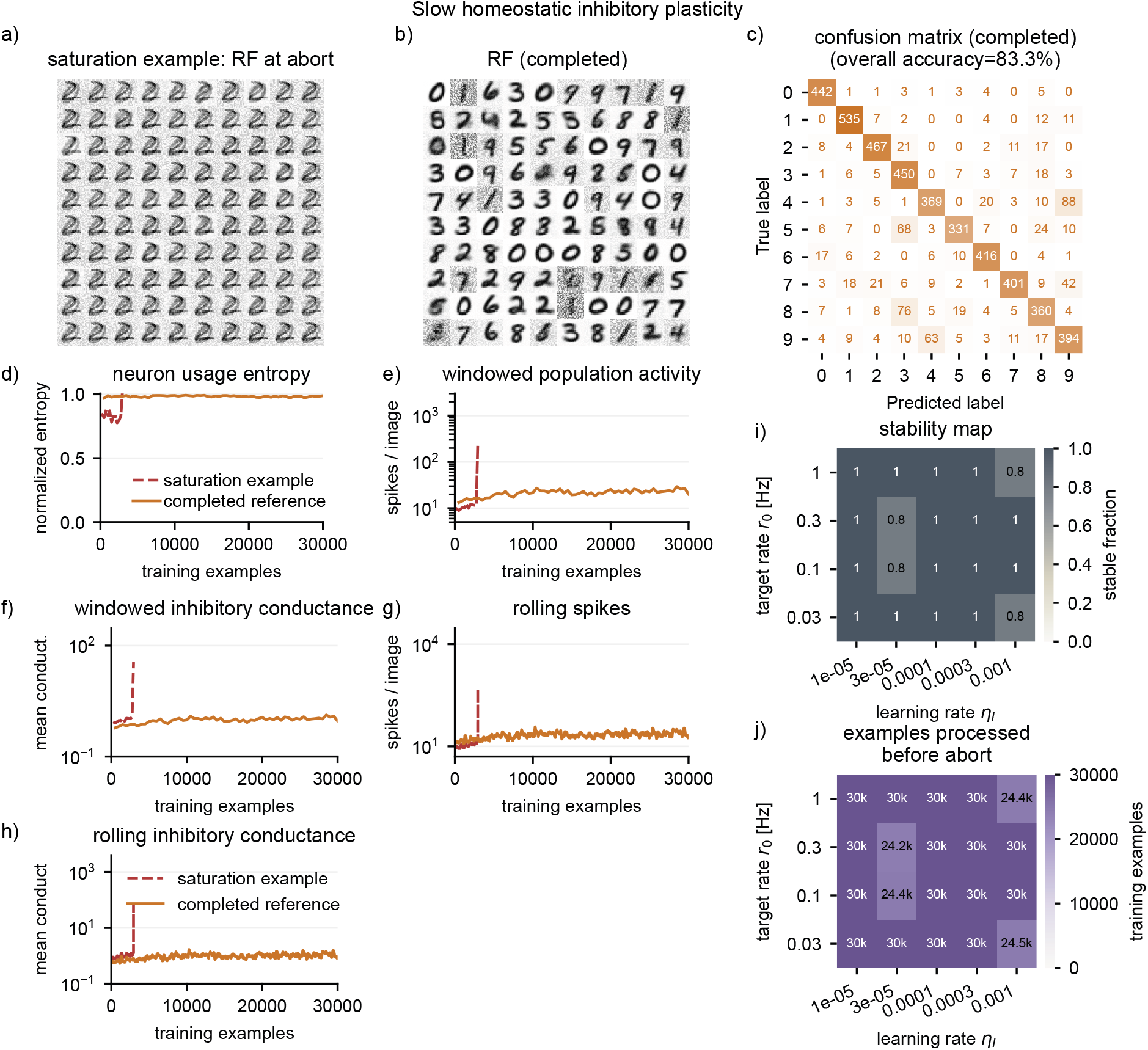
Slow homeostatic inhibitory plasticity preserves the reference regime in completed runs but can enter saturation under stress. (a,b) Input-to-excitatory receptive fields from an early saturation example at abort and from a completed reference-regime run. (c) Post hoc readout confusion matrix for the completed run; no failure confusion matrix is shown because the saturation example did not complete the training and testing workflow. (d–f) Window-based neuron-usage entropy, excitatory population activity, and inhibitory conductance for the saturation and completed examples. (g,h) Rolling single-example spike-count and inhibitory-conductance trajectories. The saturation example comes from an earlier short stress run and is used only as a diagnostic illustration of the failure mode. (i,j) Slow-homeostatic stability and processed-example maps across inhibitory learning rate *η*_*I*_ and target rate *r*_0_.

### Budget-constrained homeostatic inhibition under matched and mismatched budgets

Finally, we tested whether explicitly conserving the total inhibitory input could make slow homeostatic inhibition more controlled. The budget-control argument formalized in Eqs. (14) to (16) separates two effects that are otherwise coupled: local inhibitory updates can redistribute synaptic weights and change the total inhibitory budget received by an excitatory neuron, while column-wise normalization enforces a chosen budget *B*_*I*_. This led to the prediction that budget-constrained plasticity should preserve competition only when *B*_*I*_ is chosen from a stable fixed-inhibition regime. The final inhibitory weights in Fig. 3b show the consequence of choosing the wrong budget. When the budget-constrained rule used the mismatched budget *B*_*I*_ = 7980, its mean inhibitory weight was driven to approximately 19.95, twice the inhibitory strength of the stable fixed baseline. The corresponding rule stability in Fig. 3a shows that this mismatched budget produced runaway activity in all five replicate initializations, while the matched-budget version completed all five. The supporting budget analysis shows the same point from the fixed-regime side (Fig. S2a,b). Here, at input gain *g* = 2.0, the budget corresponding to 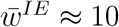 was stable in all fixed-inhibition replicates, while larger budgets were less stable and associated with elevated activity.

When the budget was matched to the stable fixed-inhibition regime (*B*_*I*_ = 3990), the budget-constrained rule reached 83.5 ± 0.6% test accuracy, preserved population activity close to fixed inhibition (21.53 ± 0.07 spikes per image), and followed the same inhibitory-conductance trajectory as the other stable matched-budget conditions (Fig. 3c–e). In Fig. 3f, its final mean inhibitory weight remained at the expected budget-implied value (9.975), while the mismatched control enforced the larger mean weight near 20 (Fig. 3b; Fig. S2c,d). This contrast becomes even more explicit in the rule-specific diagnostics of Fig. 6. The mismatched budget produced repetitive receptive fields at abort (Fig. 6a), while the matched budget retained recognizable receptive fields and a structured confusion matrix (Fig. 6b,c). The matched trajectory preserved high neuron-usage entropy and stable activity/conductance traces, while the mismatched trajectory left the reference regime and entered runaway activity (Fig. 6d–h). The stability map in Fig. 6i shows that this dependence on *B*_*I*_ generalized beyond the single matched/mismatched comparison. All runs with *B*_*I*_ = 1995 completed training, 24*/*25 runs with the matched budget *B*_*I*_ = 3990 completed training, intermediate budgets at *B*_*I*_ = 5985 were mixed, and no runs with *B*_*I*_ = 7980 completed training. The processed-example map in Fig. 6j further shows that the highest budget failed early, after only about 2700 examples on average across learning rates. The mechanistic diagnostics show why this happened. Matched budget-constrained inhibition redistributed individual weights while keeping every excitatory neuron’s total inhibitory budget at *B*_*I*_ = 3990, whereas the mismatched rule conserved the wrong budget at *B*_*I*_ = 7980 and failed (Fig. 5a–d). The correlation-based diagnostics show that this matched redistribution was not simply residual noise around a fixed matrix. Like the other plastic rules, budget-constrained inhibition produced structured inhibitory input profiles that differed from fixed inhibition but remained aligned with the learned feedforward organization (Fig. 5i–l; Fig. S3). These results show that normalization alone is not a generic stabilizer. The conserved inhibitory budget must match a stable competitive operating regime.

**Figure 5:**
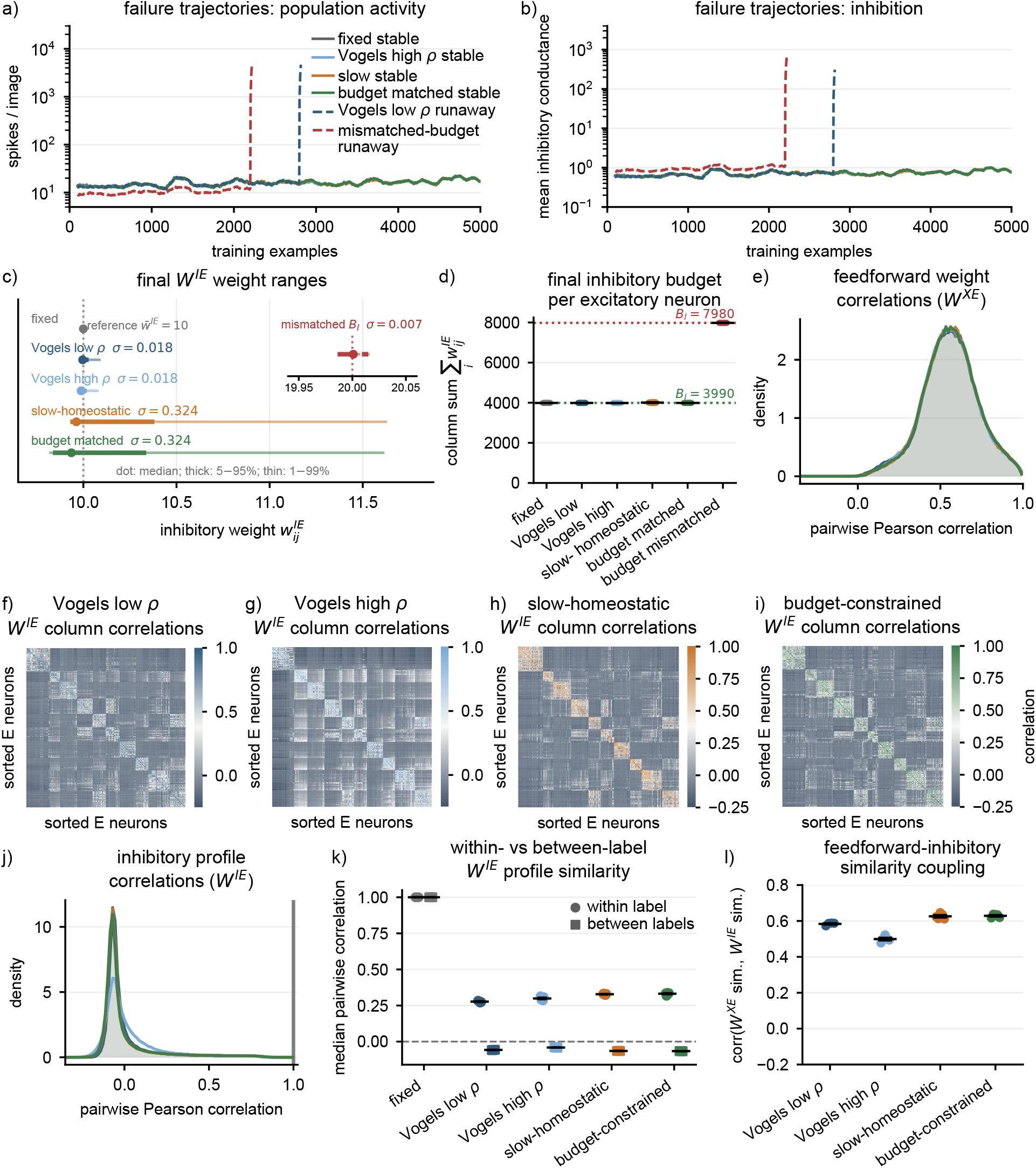
Mechanistic diagnostics distinguish stable replacement from failure. (a,b) Representative trajectories for stable matched-reference rules and two failed runs. The stable fixed, Vogels high-*ρ*, slow-homeostatic, and matched-budget runs occupied the same low-activity band, whereas a Vogels low-*ρ* run and a mismatched-budget run later showed runaway population activity and rapidly increasing inhibitory conductance. (c) Final off-diagonal inhibitory weight ranges show median, 5–95%, and 1–99% ranges for stable matched-reference rules; the inset shows the mismatched-budget control on its own scale. Slow-homeostatic and matched budget-constrained inhibition redistributed local inhibitory weights more strongly than fixed or Vogels-derived inhibition. (d) Final column-wise inhibitory budgets show the distinction between unconstrained redistribution and explicit budget conservation. The matched budget-constrained rule preserved *B*_*I*_ = 3990, while the mismatched-budget control conserved *B*_*I*_ = 7980 and failed. (e) Pairwise correlations between feedforward *W*^*XE*^ columns were similar across methods, indicating comparable receptive-field organization. (f–i) Representative inhibitory-profile correlation matrices for completed plastic-inhibition runs, with excitatory neurons sorted by post hoc digit assignment. (j) Pairwise correlations between inhibitory *W*^*IE*^ columns show the shift from uniform fixed inhibition toward redistributed plastic inhibitory profiles. (k) Within-label and between-label inhibitory-profile correlations show that plastic inhibition produced label-structured inhibitory organization. (l) Feedforward–inhibitory similarity coupling shows that excitatory neurons with more similar *W*^*XE*^ profiles tended to receive more similar *W*^*IE*^ profiles.

**Figure 6:**
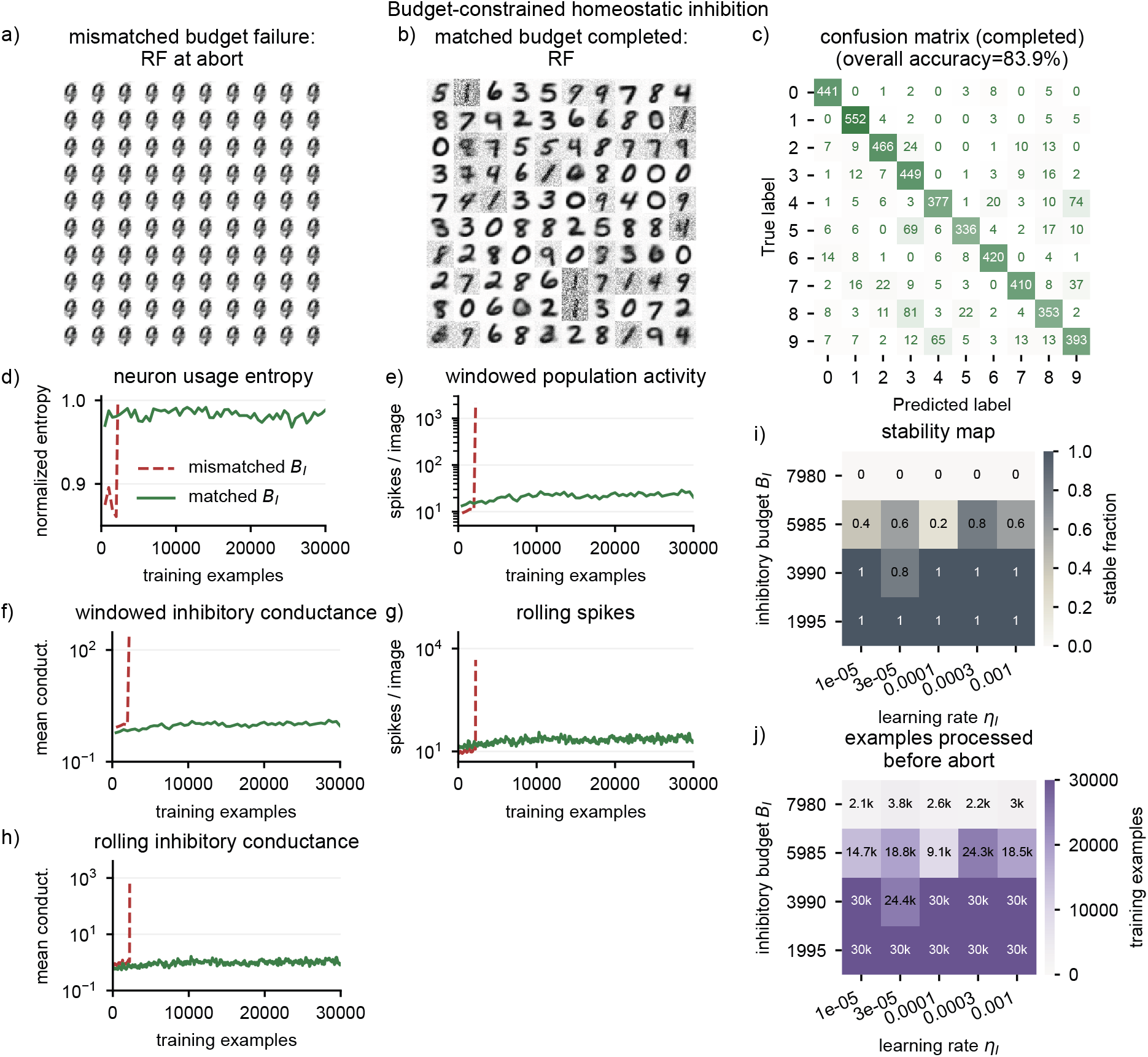
Budget-constrained homeostatic inhibition depends on matching the conserved inhibitory budget to the stable regime. (a,b) Input-to-excitatory receptive fields from the mismatched-budget run at abort and from the matched-budget run after completion. (c) Post hoc readout confusion matrix for the matched-budget run; no mismatch confusion matrix is shown because the mismatched-budget run aborted before a valid test readout. (d–f) Window-based neuron-usage entropy, excitatory population activity, and inhibitory conductance for the mismatched and matched examples. (g,h) Rolling single-example spike-count and inhibitory-conductance trajectories. (i,j) Budget-constrained stability and processed-example maps across inhibitory learning rate *η*_*I*_ and inhibitory budget *B*_*I*_.

## Discussion

We asked whether fixed lateral inhibition in a competitive STDP network can be replaced by inhibitory plasticity, motivated by the fact that biological inhibitory synapses are themselves adaptive rather than fixed [10, 14, 15, 16]. The results support a conditional answer. Inhibitory plasticity can be stable, and the Vogels-derived inhibitory STDP rule was viable across most tested parameter combinations in the selected reference regime. However, stability was not a property of the local rule alone. It depended on the operating regime of the competitive network and, for budget-constrained rules, on the inhibitory budget that defines total competition strength.

This conclusion refines the usual intuition behind homeostatic inhibitory plasticity. The inhibitory STDP rule introduced by Vogels et al. [12] is a powerful local mechanism for balancing excitation and inhibition in recurrent networks, and experimental work has shown that inhibitory spike-timing rules can interact with excitatory plasticity to shape cortical responses [13, 14]. Computational studies have extended this view by showing that inhibitory plasticity can support receptive-field formation, decorrelation, stimulus-specific competition, and stable excitatory learning when it is matched to the surrounding circuit and plasticity rules [9, 22, 23, 24]. However, inhibitory plasticity is codependent with the circuit in which it operates [15, 16]. A Diehl–Cook-style competitive architecture is not a generic balanced recurrent circuit. It includes adaptive thresholds, normalized feedforward excitatory weights, a separate inhibitory population, and stimulus-driven competition [7]. These additional stabilizing and competitive mechanisms change the dynamical context in which inhibitory plasticity acts. The stability map shows that Vogels-derived inhibitory STDP can be robust in many conditions, but also that a small number of initialization- and parameter-specific failures remain even in an otherwise stable operating regime.

The slow homeostatic rule occupies an intermediate position between spike-timing-based iSTDP and explicit budget conservation. Removing fast coincidence potentiation was sufficient to preserve the reference regime in the matched comparison, and the expanded parameter map showed that this rule was broadly stable across the tested learning rates and target rates. At the same time, without a column-wise constraint, the slow rule allowed modest variation in the total inhibition received by different excitatory neurons and could enter saturation under stress. This behavior echoes a long-standing theme in plasticity theory, where correlation-sensitive synaptic updates can create useful selectivity, but their long-term outcome is determined by the stabilizing constraints placed around them [26, 33, 34]. It also resonates with the temporal-paradox argument that slow homeostatic mechanisms are not automatically sufficient to stabilize fast Hebbian instabilities, and with recurrent-network models in which coordinated excitatory and inhibitory homeostatic rules are required to reach a desired dynamical regime [30, 35]. It therefore motivates treating the total inhibitory input as a control variable rather than as an incidental by-product of local plasticity.

The budget-constrained slow homeostatic rule makes this control variable explicit. Column-wise normalization of inhibitory weights is analogous in spirit to the feedforward weight normalization used to maintain competition among excitatory receptive fields, to classical normalized Hebbian learning rules, to biologically motivated competition for finite synaptic building blocks, and to synapse-type-specific resource competition in recurrent excitatory–inhibitory networks [32, 33, 36, 31]. The normalization target is in particula important as it fixes the total competitive drive, not merely the numerical range of individual synapses. Across the budget-constrained map, low and matched budgets were stable or nearly stable across learning rates, intermediate budgets produced mixed outcomes, and the highest budget failed in every replicate. Thus, the relevant object is not merely a plasticity rule, but a plasticity rule embedded in a constrained dynamical regime.

Biologically, this result is consistent with the broader idea that stable network learning depends on multiple interacting homeostatic processes rather than on any single local rule [28, 29, 27, 17, 30]. The inhibitory budget in the budget-constrained rule is not meant as a direct molecular mechanism. Rather, it formalizes a computational constraint, where each excitatory neuron should receive a conserved total amount of inhibitory competition while local inhibitory synapses redistribute that budget according to activity. Such a constraint could be implemented biologically by slower cell-wide or dendrite-wide homeostatic processes that interact with synapse-specific inhibitory plasticity. Models of synaptic resource competition show how local plasticity can be shaped by finite pools of synaptic building blocks, providing one mechanistic precedent for treating normalization-like constraints as biologically meaningful rather than purely numerical [36]. This is compatible with the view that inhibitory synapses contribute not only fast feedback but also slower circuit-specific regulation of excitation–inhibition balance [10, 14, 15, 16].

The computational implication is also broader than the present MNIST implementation. Competitive STDP networks, visual feature-learning models, and cortical microcircuit models all rely on a similar tension between selective Hebbian amplification and mechanisms that prevent one assembly or response pattern from dominating the network [4, 5, 37, 9]. Recent work on learned excitatory–inhibitory assemblies similarly emphasizes that inhibitory plasticity can structure stimulus-specific competition and memory-related inhibitory assemblies, not only regulate mean activity [23, 21, 25]. Eckmann, Young, and Gjorgjieva recently addressed this issue at a more general level by showing that synapse-type-specific competition for finite excitatory and inhibitory resources can organize functional recurrent rate networks with balanced receptive fields, decorrelation, response normalization, and surround suppression [31]. Our results are complementary rather than redundant. We do not model the developmental self-organization of fully plastic recurrent circuits or the disinhibitory recall of embedded engrams. Instead, we translate the broader idea of synaptic resource constraints into a practical replacement test for a spiking Diehl–Cook-style STDP network: local inhibitory weights may adapt, but the total inhibitory drive must remain matched to the stable operating regime.

The weight-correlation analysis clarifies what “preserving the regime” means mechanistically. At the feedforward level, pairwise *W*^*XE*^ correlations were almost unchanged across successful methods, consistent with the similar receptive fields and readout performance. At the inhibitory level, however, the plastic rules were not equivalent to fixed inhibition. Fixed inhibition produced nearly identical inhibitory input profiles across excitatory neurons, whereas Vogels-derived iSTDP, slow homeostatic inhibition, and budget-constrained homeostasis redistributed inhibitory profiles into label-structured patterns. Excitatory neurons assigned to the same digit tended to receive similar inhibitory profiles, while neurons assigned to different digits often received weakly anticorrelated or complementary inhibitory profiles. These negative profile correlations do not mean that inhibition became excitatory. Instead they mean that different excitatory neurons were inhibited by different relative combinations of inhibitory partners. Thus, successful plastic-inhibition conditions can look similar at the output level while differing substantially in their hidden inhibitory organization. This observation is conceptually aligned with recent work in recurrent memory networks, where inhibitory plasticity shapes structured inhibitory assemblies rather than merely stabilizing average activity [25]. In our model, however, the structure is read out as label-aligned inhibitory profiles in a competitive representation-learning task rather than as an explicitly embedded inhgram supporting recall. The output-level similarity is therefore not a failure to change the model. It is evidence that adaptive inhibition preserved the functional regime that fixed inhibition previously imposed, while reorganizing the internal inhibitory scaffold within that regime.

This regime-level view is also consistent with recent automated searches over multi-type plasticity rules, which found diverse sets of co-active local rules that stabilize recurrent spiking networks and support memory functions, while individual rules often failed when considered in isolation [38]. That work explores a broad discovery problem over coupled plasticity rules across multiple recurrent synapse classes. In contrast, the present study isolates a narrower replacement problem in a competitive STDP architecture and identifies inhibitory budget matching as one concrete constraint that preserves the fixed-inhibition operating regime.

The present experiments deliberately prioritize stability analysis over exhaustive task optimization, which defines the scope of the conclusions. We used five replicate initializations, 30,000 training examples, and 5000 test examples, thus, a scale sufficient to characterize replicate stability regimes, but smaller than exhaustive MNIST benchmarking. The aim was to analyze stability mechanisms rather than to optimize final classification performance. Accordingly, differences from the strongest reported Diehl–Cook-style MNIST accuracies may reflect the Euler implementation, finite replicate-sweep protocol, reduced benchmark subset, and the absence of task-specific parameter optimization. The same considerations likely explain why some excitatory receptive fields remained noisy or weakly specialized. The longer fixed-inhibition diagnostic improved neither accuracy nor receptive-field organization enough to suggest that the core conclusions would change simply by extending the baseline run. The network implementation also remains close to the Diehl–Cook architecture and does not test deeper spiking hierarchies, continuous naturalistic inputs, behavioral time series, or plasticity within the inhibitory population itself. Finally, the budget-constrained slow homeostatic rule is only one point in a larger space of constrained inhibitory plasticity mechanisms. Future work should test whether the same budget principle transfers to nonstationary inputs, larger networks, more biologically detailed inhibitory populations, and co-active I-to-E/I-to-I plasticity regimes in which inhibition becomes an assembly-specific substrate for disinhibitory computation [25].

Taken together, these results support both a practical and theoretical conclusion. In competitive STDP networks, replacing fixed inhibition with inhibitory plasticity is not a plug-in substitution. Local inhibitory learning rules must be evaluated together with the competitive operating regime and the total inhibitory budget they imply. Budget-matched inhibitory plasticity offers a simple route toward plastic yet stable competition, while the failures of mismatched budgets emphasize that homeostatic plasticity can stabilize only the regime it is constrained to preserve. In this sense, inhibitory budget matching instantiates a broader resource-constraint principle in a concrete spiking STDP setting: it separates local redistribution of inhibitory weights from changes in the total competition strength that defines the functional regime. This provides a concrete starting point for more general theories of inhibitory learning in competitive spiking networks.

## Data and code availability

All simulations were generated with the accompanying Euler-based spiking-network implementation. The source code, analysis scripts, and HPC workflow are available on GitHub at https://github.com/FabrizioMusacchio/inhibitory_budget_stdp and archived on Zenodo [39]: https://doi.org/10.5281/zenodo.22976646. The simulation outputs required to reproduce the data-driven figure panels (long_30k_sweeps_manifest_HPC/, slow_budget_maps_30k_manifest_HPC/, and long_visualization_60k_2ep/) are archived separately on Zenodo [40]: https://doi.org/10.5281/zenodo.22976539.

## Acknowledgements

The author thanks colleagues at DZNE for informal discussions and feedback on the project.

## Author contributions

F.M. conceived the study, developed the model and analysis code, performed the simulations, analyzed the data, generated the figures, and wrote the manuscript. M.F. supervised the project, acquired funding, and reviewed the manuscript.

## Use of AI-assisted tools

The authors used OpenAI ChatGPT, GPT-5, to assist with language editing and wording refinement. The tools were not used to generate primary research data, perform unsupervised interpretation of results, or make authorship-level scientific decisions. All AI-assisted text changes were reviewed, verified, and edited by the authors, who take full responsibility for the accuracy, integrity, and content of the submitted article.

## Supplementary Information

**Figure S1:**
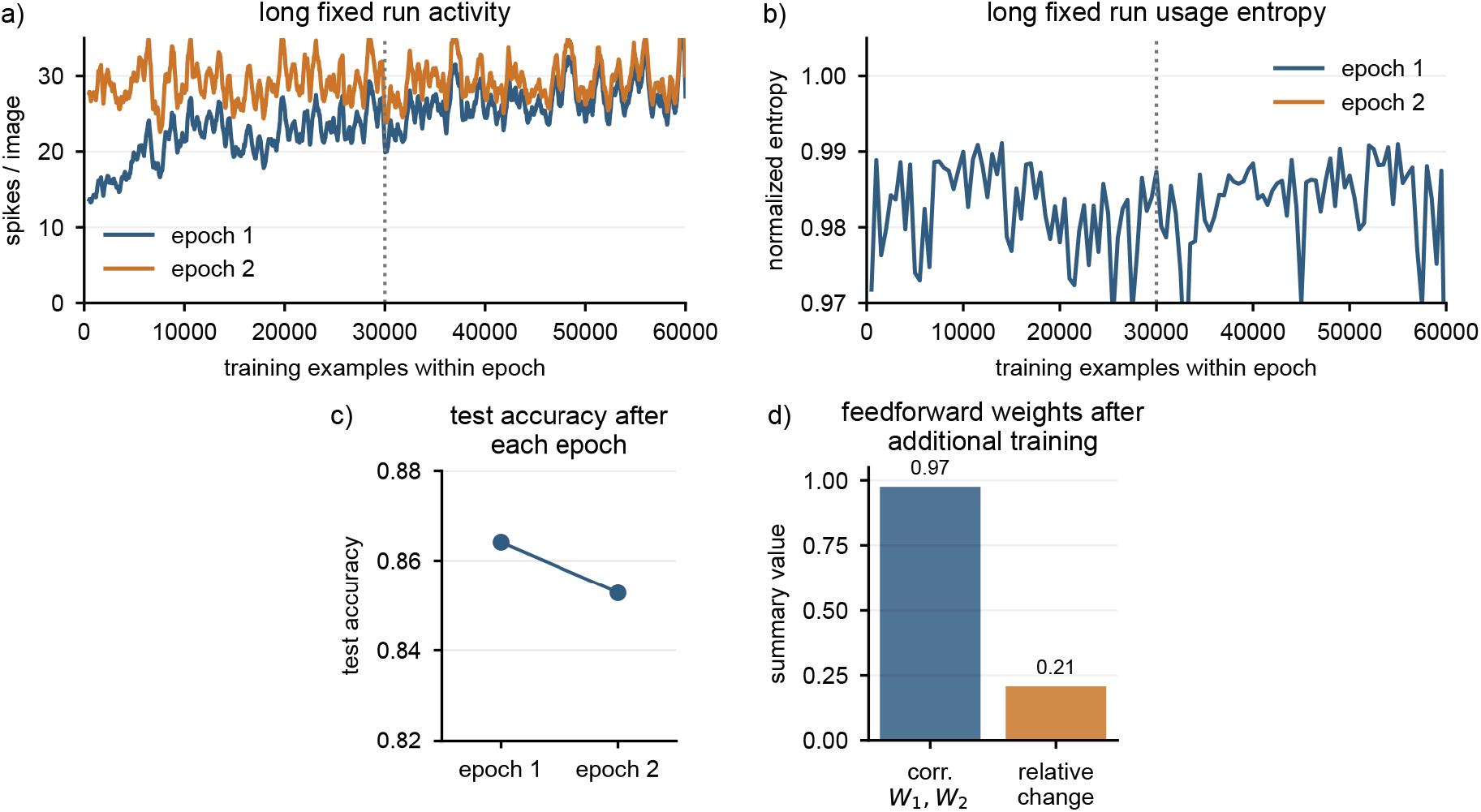
Long fixed-inhibition diagnostic run used to choose a tractable sweep protocol. A fixed-inhibition run with 60,000 training examples per epoch and two epochs was used to assess whether the shorter replicate sweeps captured stable representation dynamics. (a) Rolling mean excitatory spikes per image remained in a stable range during both epochs; the dotted line marks 30,000 examples. (b) Neuron usage entropy was already high during the first epoch and remained high in the second epoch. (c) Test accuracy did not improve after the second epoch (86.4% after epoch 1 and 85.3% after epoch 2). (d) Input-to-excitatory weights remained strongly correlated between the end of epoch 1 and the end of epoch 2. These diagnostics motivated the 30,000-training-example, 5000-test-example protocol used for the replicate regime sweeps.

**Figure S2:**
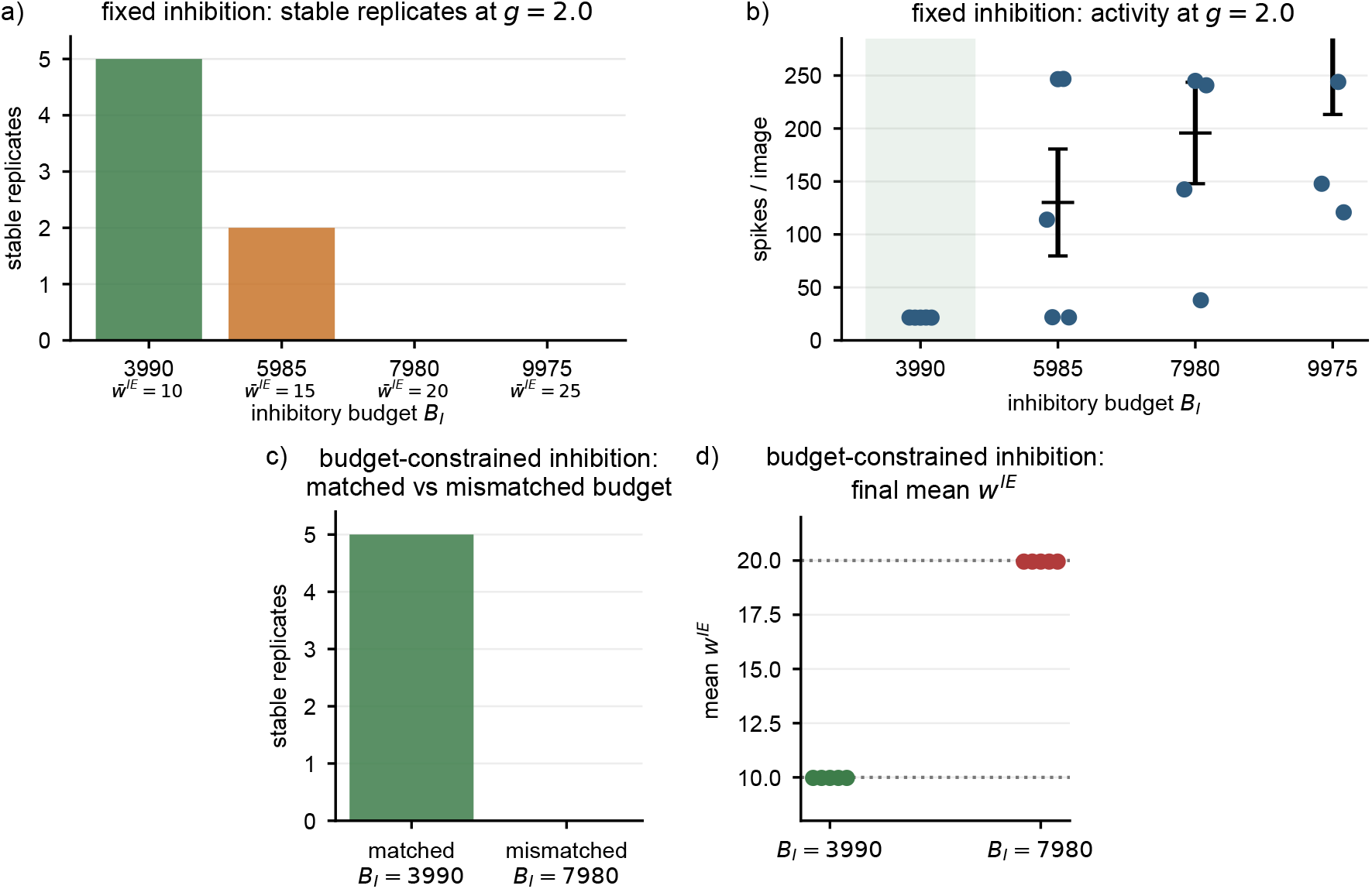
Evidence for the matched and mismatched inhibitory budgets. Self-inhibition is excluded in the matched inhibitory–excitatory populations, so each excitatory neuron receives inhibition from *N*_*I*_ − 1 = 399 inhibitory neurons and 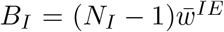. (a) In the fixed-inhibition sweep at input gain *g* = 2.0, the budget corresponding to 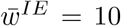 (*B*_*I*_ = 3990) completed all five replicate initializations, while larger fixed-inhibition budgets were less stable. (b) The same sweep showed that larger budgets were associated with much higher excitatory activity among completed or aborted runs. (c) For the budget-constrained slow-homeostatic rule, the budget matched to the fixed stable regime (*B*_*I*_ = 3990) was stable in all five replicate initializations, while the mismatched budget (*B*_*I*_ = 7980) was unstable in all five replicate initializations. (d) Final inhibitory weights confirmed that the matched and mismatched budget-constrained rules enforced mean inhibitory magnitudes near 10 and 20, respectively.

**Figure S3:**
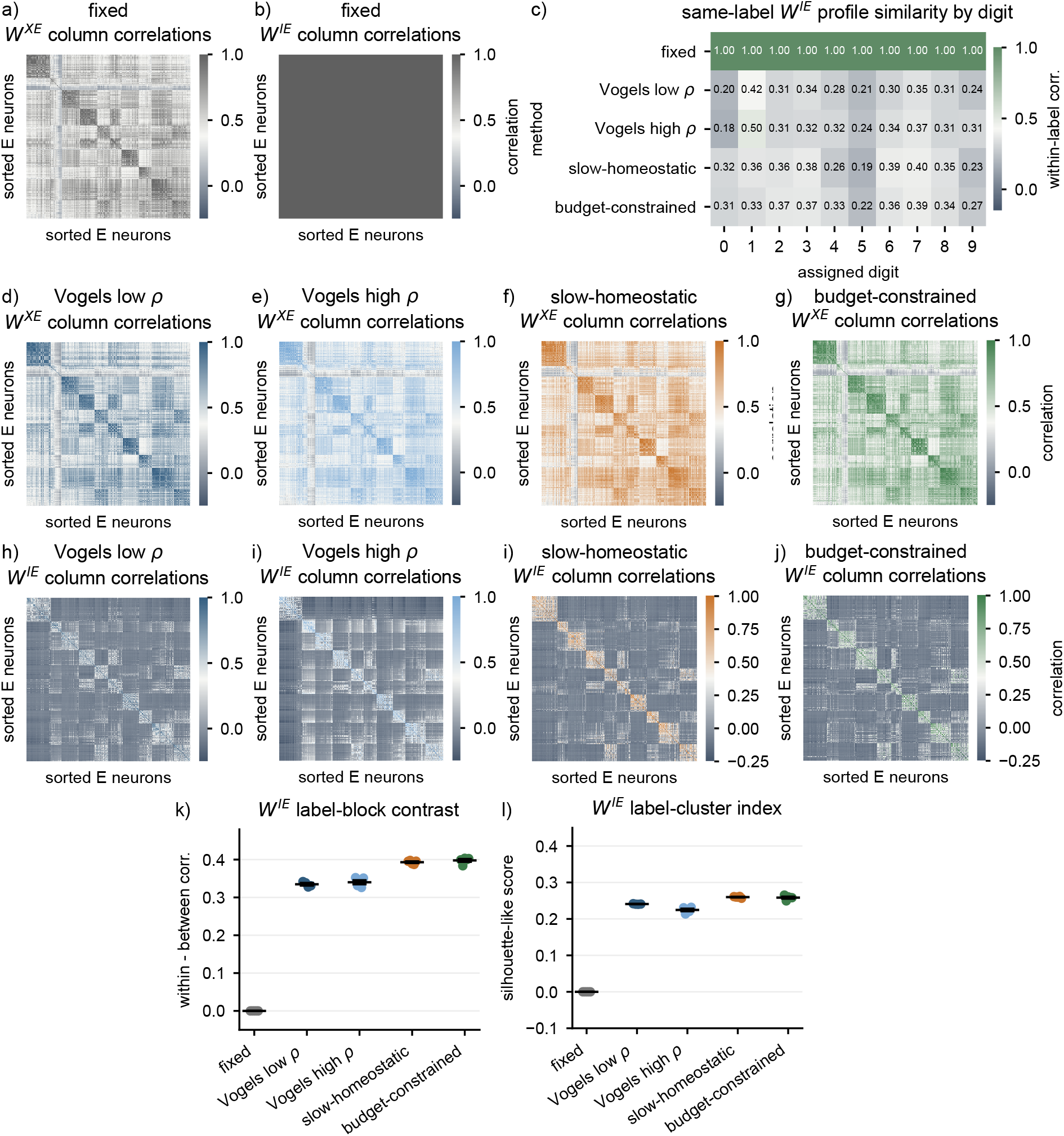
Weight-correlation diagnostics for feedforward and inhibitory profiles. Pairwise Pearson correlations were computed between excitatory-neuron weight columns after sorting excitatory neurons by their post hoc digit assignment. (a,b) Fixed-inhibition reference: feedforward *W*^*XE*^ columns show structured receptive-field similarity, whereas inhibitory *W*^*IE*^ columns are nearly perfectly correlated because fixed inhibition gives each excitatory neuron the same inhibitory profile. (c) Same-label *W*^*IE*^ profile similarity by assigned digit. Plastic rules produced positive within-label correlations, with digit-to-digit variation in how strongly inhibitory profiles clustered. (d–g) Feedforward *W*^*XE*^ correlation matrices for the plastic-inhibition methods show that the learned feedforward organization remained broadly similar across methods. (h–j) Corresponding inhibitory *W*^*IE*^ correlation matrices show that plastic inhibition redistributed the hidden inhibitory profiles into non-uniform, label-structured patterns. (k) Label-block contrast, computed as within-label minus between-label inhibitory-profile correlation. (l) Silhouette-like label-cluster index for inhibitory-profile organization. Together, these diagnostics show that successful plastic inhibition preserved the feedforward representation while reorganizing the hidden inhibitory scaffold.

